# Regulation of antimycin biosynthesis is controlled by the ClpXP protease

**DOI:** 10.1101/576462

**Authors:** Bohdan Bilyk, Sora Kim, Asif Fazal, Tania A. Baker, Ryan F. Seipke

**Affiliations:** Astbury Centre for Structural Molecular Biology, Faculty of Biological Sciences, University of Leeds, Leeds, UK; Department of Biology, Massachusetts Institute of Technology, Cambridge, MA, USA; Howard Hughes Medical Institute, Chevy Chase, MD, USA

**Keywords:** *Streptomyces*, regulation of secondary metabolism, antimycin, ECF sigma factors, proteolysis, ClpXP

## Abstract

The survival of any microbe relies upon its ability to respond to environmental change. Use of Extra Cytoplasmic Function (ECF) RNA polymerase sigma (σ) factors is a major strategy enabling dynamic responses to extracellular signals. *Streptomyces* species harbor a large number of ECF σ factors; nearly all of which regulate genes required for morphological differentiation and/or response to environmental stress, except for σ^AntA^, which regulates starter-unit biosynthesis in the production of antimycin, an anticancer compound. Unlike a canonical ECF σ factor, whose activity is regulated by a cognate anti-σ factor, σ^AntA^ is an orphan, raising intriguing questions about how its activity may be controlled. Here, we reconstitute *in vitro* ClpXP proteolysis of σ^AntA^, but not a variant lacking a C-terminal di-alanine motif. Furthermore, we show that the abundance of σ^AntA^ *in vivo* is enhanced by removal of the ClpXP recognition sequence, and that levels of the protein rise when cellular ClpXP protease activity is abolished. These data establish direct proteolysis as an alternative and thus far unique control strategy for an ECF RNA polymerase σ factor and expands the paradigmatic understanding of microbial signal transduction regulation.

**Importance:** Natural products produced by *Streptomyces* species underpin many industrially- and medically-important compounds. However, the majority of the ~30 biosynthetic pathways harboured by an average species are not expressed in the laboratory. This undiscovered biochemical diversity is believed to comprise an untapped resource for natural products drug discovery. A major roadblock preventing the exploitation of unexpressed biosynthetic pathways is a lack of insight into their regulation and limited technology for activating their expression. Our findings reveal that the abundance of σ^AntA^, which is the cluster-situated regulator of antimycin biosynthesis, is controlled by the ClpXP protease. These data link proteolysis to the regulation of natural product biosynthesis for the first time and we anticipate that this will emerge as a major strategy by which actinobacteria regulate production of their natural products. Further study of this process will advance understanding of how expression of secondary metabolism is controlled and will aid pursuit of activating unexpressed biosynthetic pathways.

## Introduction

The survival of any organism relies upon its ability to respond to environmental change. This feature is especially true of bacteria, which often live in hostile and fluctuating environments. *Streptomyces* bacteria thrive in soils. The success of this genus of filamentous, sporulating bacteria is linked to their complex lifecycle and keen ability to sense and respond to its surroundings. Notably, a multitude of bioactive secondary or specialized metabolites are produced in response to environmental cues (1). More than half of all small molecule therapeutics critical for human health and wellbeing are derived from or inspired by *Streptomyces* natural products (2).

*Streptomyces* species typically harbour a large number of biosynthetic pathways, but only a few of them are expressed under common laboratory conditions. The biochemical diversity encoded by these silent pathways is a tremendous untapped resource for discovery of new antibacterial agents and other therapeutics. All available data indicates that the production of natural products is controlled predominantly at the level of transcription. Although there are complex regulatory cascades that tightly control expression of biosynthetic genes, they are ultimately activated, repressed or de-repressed by so-called cluster-situated regulators—regulatory protein(s) encoded within the biosynthetic gene cluster (BGC) (3, 4). Major roadblocks preventing the exploitation of silent biosynthetic pathways are a lack of insight into their regulation and limited technology for activating their expression.

Antimycins have been known for 70 years and are the founding member of a large class of natural products widely produced by *Streptomyces* species (5–6). Recently, antimycins were shown to be potent and selective inhibitors of the mitochondrial Bcl-2/Bcl-X_L_-related antiapoptotic proteins that are overproduced by cancer cells and confer resistance to chemotherapeutic agents whose mode of action is activation of apoptosis (7). The ~25 kb antimycin (*ant*) BGC harboured by *S. albus* is composed of 15 genes organised into four polycistronic operons: *antBA, antCDE*, *antGF* and *antHIJKLMNO* (Fig. 1) (8, 9). The regulation of the *ant* BGC is unusual compared to other secondary metabolites. Its expression is regulated by FscRI, a cluster-situated LuxR-family regulator of candicidin biosynthesis; FscRI activates expression of *antBA* and *antCDE* (10). Importantly, *antA* is a cluster-situated regulator that encodes an Extra Cytoplasmic Function (ECF) RNA polymerase σ factor (σ^AntA^) that activates expression of the remaining operons: *antGF* and *antHIJKLMNO* (Fig. 1) (9).

**Fig. 1.**
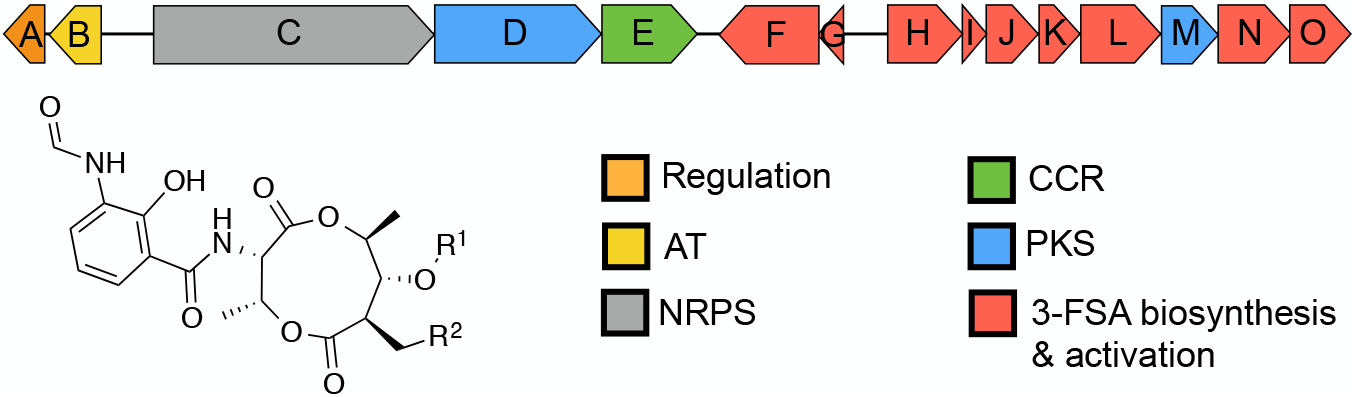
Schematic representation of the antimycin (*ant*) biosynthetic gene cluster. AT, acyltransferase; NRPS, nonribosomal peptide synthetase; PKS, polyketide synthase; CCR, crotonyl-CoA carboxylase/reductase, 3-FSA, 3-formamidosalicylate. Antimycins: Antimycin A_1_, R^1^= COCH(CH_3_)CH_2_CH_3_, R^2^= (CH_2_)_4_CH_3_; Antimycin A_2_, R^1^=COCH(CH_3_)_2_, R^2^= (CH_2_)_4_CH_3_; Antimycin A_3_, R^1^= COCH_2_CH(CH_3_)_2_, R^2^= (CH_2_)_2_CH_3_; Antimycin A_4_, R^1^= COCH(CH_3_)_2_, R^2^= (CH_2_)_2_CH_3_.

σ^AntA^, like all ECF σ factors, is similar to the housekeeping σ^70^ family, but only possesses two of the four highly characteristic sigma domains: domains σ2 and σ4; these regions of sigma factors bind the −10 and −35 promoter elements respectively, and are sufficient for recruitment of RNA polymerase (11). Genes encoding ECF σ factors are almost always co-transcribed with their cognate anti-σ factor (12). This class of anti-σ factors are transmembrane proteins that selectively bind to and inactivate a partner σ factor until its release is stimulated, usually by an exogenous signal (12, 13). After the σ factor is released, it recruits RNA polymerase to express a defined regulon that usually includes the σ factor-anti-σ factor operon itself, which thus establishes a positive auto-feedback loop in the presence of the inducing stimulus. *Streptomyces* species encode a large number of ECF σ factors (>30 per strain) and nearly all of these regulate genes required for morphological differentiation and/or response to environmental stress and are not dedicated regulators of one biosynthetic pathway. Interestingly, unlike the canonical ECF σ factors, whose activities are controlled by cognate anti-σ factors, σ^AntA^ lacks an identifiable anti-σ factor partner and as a consequence has created curiosity about how its activity is controlled.

The Clp-protease system is essential for normal bacterial proteostasis and is best characterized in *Escherichia coli* (14, 15). The Clp protease is a multi-enzyme complex composed of a barrel-shaped peptidase, ClpP and a regulatory enzyme, either ClpA or ClpX (or ClpC in some organisms). ClpA and ClpX (and ClpC) are all AAA+−family protein unfoldases that recognise an N- and/or C-terminal recognition signal (degron) and utilise ATP to unfold and translocate proteins to the peptidase chamber where they are degraded into short peptides (16). In *Streptomyces* species, the peptidase is specified by two genes instead of one and is redundantly encoded (17). The primary peptidase is encoded by *clpP1P2*, whose corresponding proteins form a complex with ClpX or ClpA to facilitate normal proteostasis; the second peptidase is encoded by *clpP3P4,* but its expression only occurs when the primary system is compromised (18, 19). The best understood degron is the SsrA tag from *E. coli* (AANDENYALAA), which is added co-translationally to polypeptides stalled on ribosomes (20, 21). The *E. coli* SsrA tag has been comprehensively studied and the C-terminal Ala-Ala-COO^−^of this motif is essential for proteolysis by ClpXP (22). Intriguingly, the C-terminus of σ^AntA^ harbours the sequence Ala-Ala-COO^−^, which previously led us to speculate that ClpXP may modulate its level/activity (9).

Here, we reconstitute ClpXP proteolysis of σ^AntA^ *in vitro* and show that it is dependent upon the C-terminal Ala-Ala. We also show that the abundance of σ^AntA^ *in vivo* is higher when Ala-Ala is changed to Asp-Asp and that abundance σ^AntA^ is elevated in the absence of genes encoding the primary peptidase, ClpP and its unfoldase, ClpX. These data establish direct proteolysis as an alternative, and thus far unique, control strategy of ECF RNA polymerase σ factors, expanding the paradigmatic understanding of microbial signal transduction regulation.

## Results and discussion

### σ^AntA^ orthologues are a new subfamily of ECF σ factor that regulate production of the antimycin biosynthetic starter unit

Since its initial discovery six years ago, more than 70 *ant* BGCs have been identified within the Actinobacteria including *Actinospica*, *Saccharopolyspora*, *Streptacidiphilus* and *Streptomyces* (5). Each of these BGCs harbours a single regulator, σ^AntA^ (53-100% shared amino acid identity across all orthologues), which lacks a cognate anti-σ factor partner (5, 9). Our previous work with *S. albus* S4 established that σ^AntA^ orthologues comprise a new subfamily of ECF σ factors (9, 23). We demonstrated σ^AntA^ is required for expression of *antGF* and *antHIJKLMNO*, which encode a standalone ketoreductase (AntM) and proteins required for the production/activation of the starter unit, 3-formamidosalicylate (3-FSA) (Fig. 1). We also mapped the transcriptional start sites and identified conserved promoter sequences for these operons in all known antimycin BGCs at the time (9). The conservation of σ^AntA^ and target promoters within *ant* BGCs from taxonomically diverse species, suggests that σ^AntA^-mediated regulation of these genes is direct. To verify this hypothesis, we performed ChIP-sequencing with a *S. albus* S4 Δ*antA* mutant complemented with an N-terminal 3xFLAG-tagged version of σ^AntA^. The number of reads that mapped to the promoters of *antGF and antHIJKLMNO* was enriched for both biological replicates of Δ*antA*/3xFLAG-*antA* compared to that of the wild-type mock-immunoprecipitated control, indicating that σ^AntA^ presumably directly activates the production of the 3-FSA starter unit during antimycin biosynthesis (Fig. 2).

**Fig. 2.**
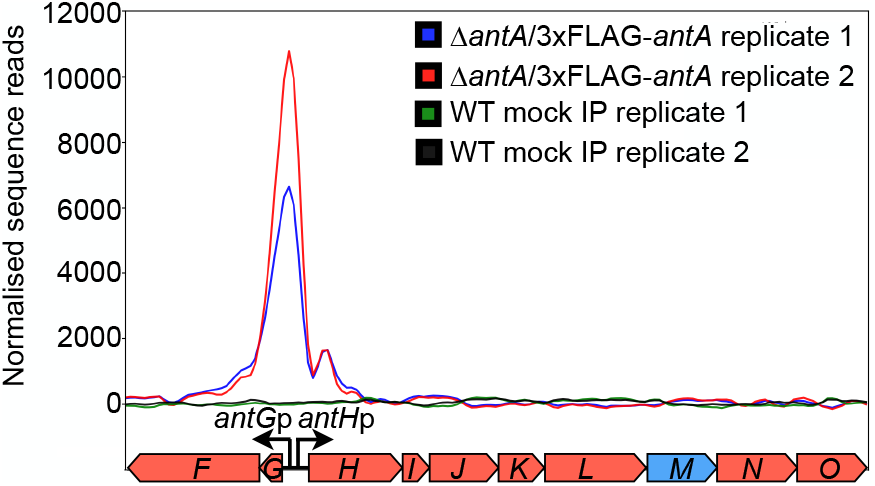
3xFLAG-σ^AntA^ binds to the *antGF* and *antHIJKLMNO* promoters *in vivo.* Shown is a graphical representation of normalised sequence reads mapped to the intergenic region of *antG-antH* (shown at bottom). The genomic coordinates depicted are nucleotides 43,148 to 51,448 of contig CADY01000091.1 of the *S. albus* S4 genome^49^. WT, wild-type; IP, immunoprecipitation.

### σ^AntA^ is degraded by the ClpXP protease *in vitro*

The activities of almost all characterized ECF σ factors are modulated by a cognate anti-σ factor, which is typically a small transmembrane protein co-encoded within the same operon. Intriguingly, σ^AntA^ lacks an anti-σ factor and is therefore an orphan, indicating a unique mechanism is likely at work to control σ^AntA^ activity. An inspection of σ^AntA^ amino acid sequences revealed a C-terminal Ala-Ala in 67 out of the 71 orthologues (Fig. S1). A C-terminal Ala-Ala is an important component of a common class of degrons for the ClpXP protease (22). This observation led us to hypothesize that the activity of σ^AntA^ could be modulated by proteolysis instead of by an anti-σ factor. To test this hypothesis, we performed *in vitro* proteolysis. Previous work indicated that *S. albus* S4 σ^AntA^ was insoluble when overproduced by *E. coli*, so we pursued the overproduction and purification of the orthologue from *Streptomyces ambofaciens* ATCC 23877, which is an experimentally demonstrated producer of antimycins (24). *S. ambofaciens* σ^AntA^ (75% shared amino acid identity with *S. albus* S4 σ^AntA^) was purified as an N-terminal (His)_6_-SUMO-fusion protein. The (His)_6_-SUMO tag increases solubility and eases purification of putative substrates, without altering recognition of C-terminal degrons by ClpXP. ClpX orthologues from *E. coli* and *S. ambofaciens* possess 60% shared amino acid identity and therefore likely recognise similar substrates for degradation. Thus, ClpXP from *E. coli* was purified (Fig. S2) and its ability to degrade (His)_6_-SUMO-σ^AntA^ was assessed. Degradation of (His)_6_-SUMO-σ^AntA^ was apparent as early as 2.5 min after addition of ATP and all of the sample was degraded by 15 min (Fig. 3). Substrates of ClpXP become resistant to proteolysis by specific alterations of the C-terminal Ala-Ala (22). Therefore, to investigate degradation specificity in the above experiment we constructed and tested a variant of *S. ambofaciens* σ^AntA^ in which the C-terminal Ala-Ala was mutated to Asp-Asp ((His)_6_-SUMO-σ^AntA-DD^). Strikingly, the Asp-Asp variant was stable against ClpXP degradation over the lifetime of the assay (Fig. 3). Thus, the degradation of (His)_6_-SUMO-σ^AntA^ and the characteristic resistance afforded by the Ala-Ala-to-Asp-Asp mutation demonstrates that σ^AntA^ is a substrate of ClpXP *in vitro*.

**Fig. 3.**
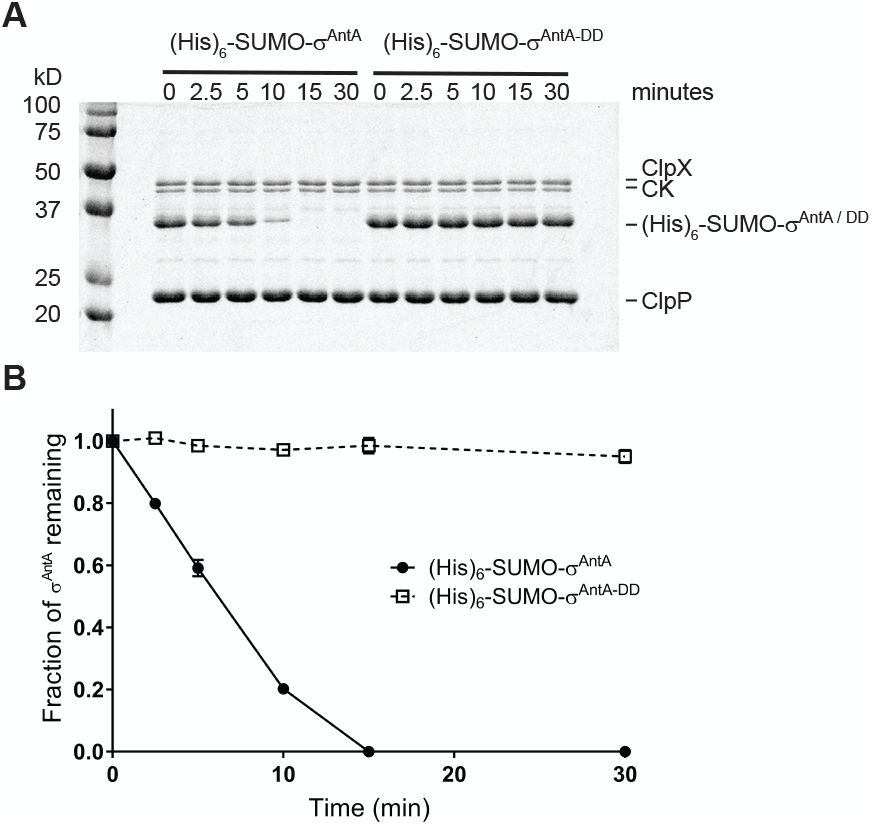
Proteolysis of *S. ambofaciens* σ^AntA^ by ClpXP in vitro. (A) SDS-PAGE analysis of proteolysis reactions containing 37 pmols (His)_6_SUMO-σ^AntA^ or (His)_6_SUMO-σ^AntA-DD^. (B) Densitometry analysis SDS-PAGE images for three independent proteolysis experiments. The mean is plotted and error bars illustrate the standard error of the mean (±1 SEM).

### σ^AntA^ is degraded by the ClpXP protease *in vivo*

To investigate if the *in vitro* degradation of σ^AntA^ demonstrated above is relevant to its regulation *in vivo*, we deleted the operon encoding the *clpX, clpP1, and clpP2* genes from *S. albus* S4. The resulting Δ*clpXclpP1clpP2* mutant underwent a normal developmental cycle, albeit sporulation was less robust, which is consistent with growth characteristics reported for mutation of equivalent genes in *S. coelicolor* (Fig. S3) (25). Next, genes encoding the 3xFLAG-σ^AntA^ or 3xFLAG-σ^AntA-DD^ fusion proteins were generated and introduced into the parental strain and the Δ*clpXclpP1clpP2* mutant so the abundance of these proteins could be assessed over a developmental time course by Western blotting with anti-FLAG antisera. This experiment was initially performed with the σ^AntA^ fusions integrated on the chromosome under control of the native promoter. However, a reliable signal could not be detected for 3xFLAG-σ^AntA^ and only a trace amount of the Asp-Asp variant was observed, presumably indicating that the cellular level of σ^AntA^ is normally low because the native promoter is relatively weak. The experiment was therefore repeated with 3xFLAG-σ^AntA^ and 3xFLAG-σ^AntA-DD^ expression driven by a stronger, constitutive promoter, *ermE\** (26). Analysis of the resulting immunoblot revealed that 3xFLAG-σ^AntA-DD^ was more abundant than 3xFLAG-σ^AntA^ in extracts prepared from vegetative mycelia of the parent and Δ*clpXclpP1clpP2* strains (Fig. 4). Strikingly, 3xFLAG-σ^AntA^ and 3xFLAG-σ^AntA-DD^ could only be detected in extracts from aerial mycelia of the Δ*clpXclpP1clpP2* strain and not the parent; the Asp-Asp variant was also present in greater relative abundance (Fig. 4), which was consistent with our previous experiments that showed the *ant* BGC is downregulated at the level of transcription upon the onset of aerial growth (9). Interestingly, the conspicuous absence of 3xFLAG-σ^AntA^ and the presence 3xFLAG-σ^AntA-DD^ in protein extracts prepared from the latest time point sampled suggests the potential involvement of an additional degradative factor(s). Taken together, these data support the hypothesis that σ^AntA^ levels, and thus its ability to activate gene expression of *antFGHIJKLMNO* is modulated by the ClpXP protease.

**Fig. 4.**
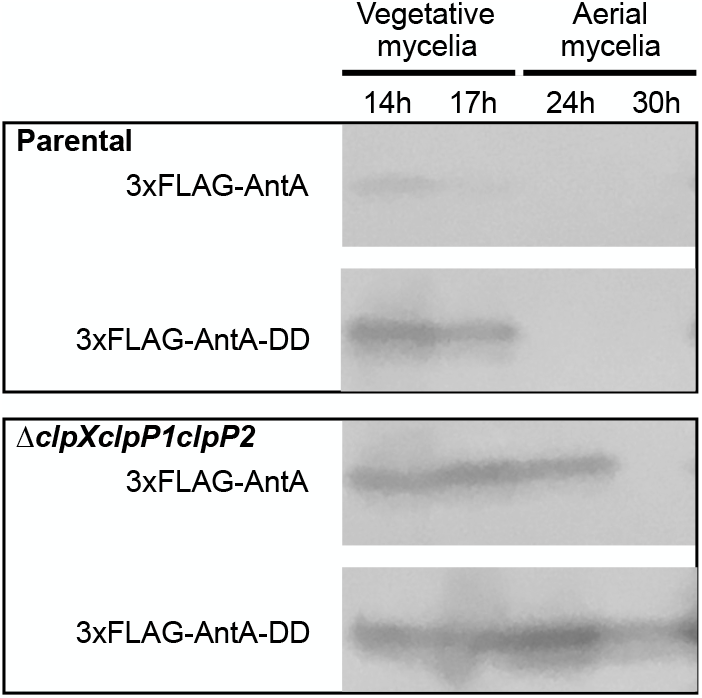
The abundance of σ^AntA^ is enhanced in the absence of the ClpXP *in vivo.* Cells from the indicated strains were cultivated over a developmental time course atop cellophane discs on agar media. Protein extracts were generated from 100 mg of either vegetative mycelia (14 and 17 hours) or aerial mycelia (24 and 30 hours). Thirty micrograms of total protein were analysed by Western blotting with anti-FLAG antisera. The images shown are derived from uncropped original images shown in Fig. S4.

**Fig. 5.**
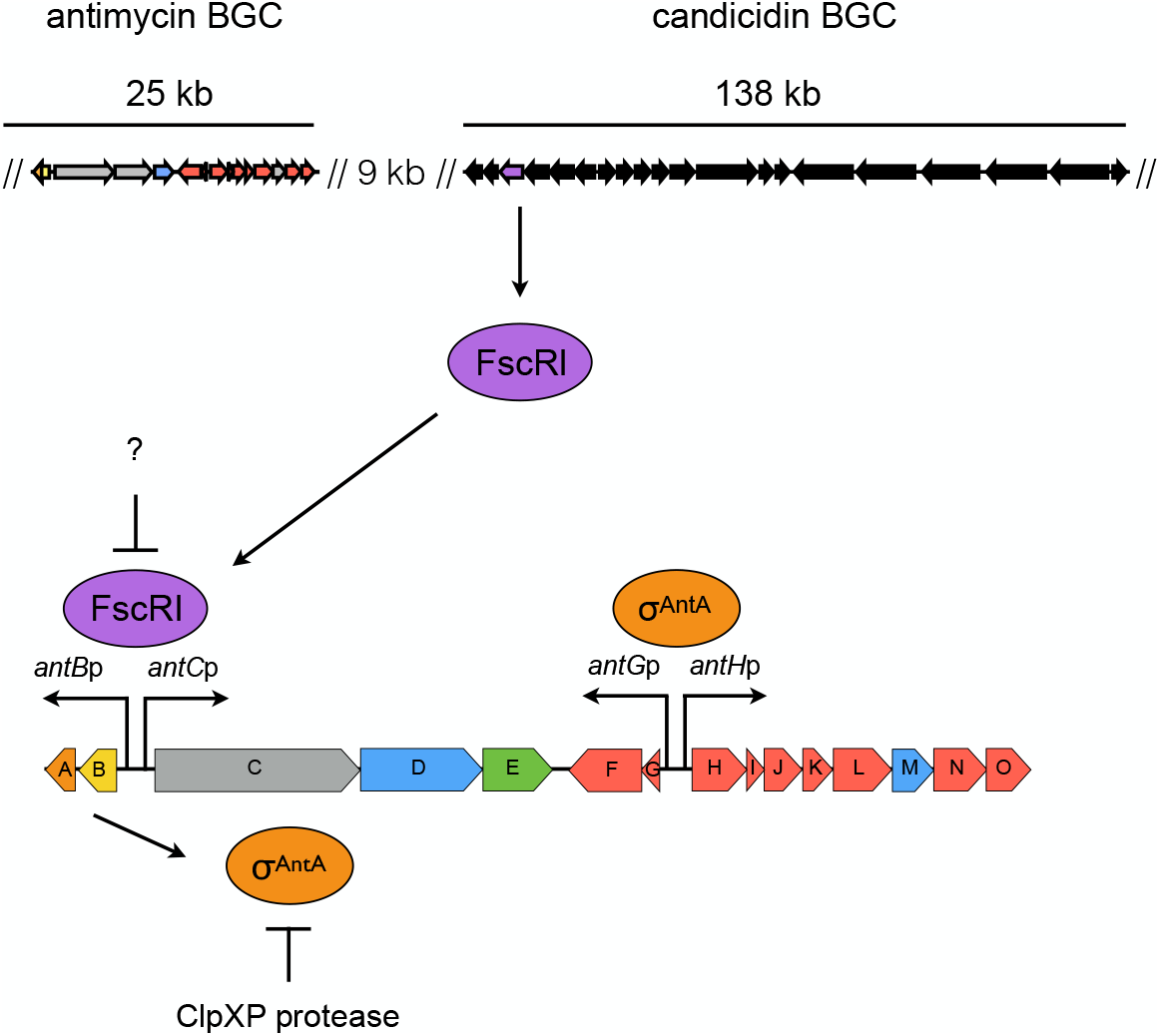
Model for the regulation of antimycin biosynthesis. The upper panel displays the relative locations of the antimycin and candicidin BGCs in the *S. albus* S4 chromosome. In the lower panel, FscRI, a LuxR-family regulator, from the candicidin BGC, activates expression of *antBA* and *antCDE*. This in turn enables direct activation of the 3-FSA biosynthetic operons (*antGF* and *antHIJKLMNO*) by σ^AntA^. The cellular level of σ^AntA^ is antagonised by ClpXP-protease system, for which it is a direct target and is ultimately responsible for clearing residual σ^AntA^ when FscRI is inactivated following the onset of differentiation.

### Antimycins are not overproduced in the absence of ClpXP

The above experiments indicate that the cellular level of σ^AntA^ is more abundant in the absence of the ClpXP protease. In order to determine if an increased level of this transcription factor ultimately influenced the final production titre of antimycins, we used LC-HRMS to assess the abundance of antimycins in chemical extracts generated from the Δ*clpXclpP1clpP2* and parental strains grown atop a cellophane disk on MS agar in triplicate. The extracted ion chromatograms representing antimycin A_1_, A_2_ A_3_ and A_4_ were used to determine the peak area for each compound, which was subsequently normalised based on the wet mycelial weight of the sample. Interestingly, the results indicated that total antimycin production by the Δ*clpXclpP1clpP2* mutant (15.57 AU ± 2.86) and parental strain (16.59 AU ± 1.12) was not statistically significantly different (*P* value 0.59) Table S1. This result is consistent with a previous experiment where overexpression of *antA* did not increase the titre of antimycins, because it only results in overexpression of *antGF* and *antHIJKLMNO* (genes encoding the production of the AntG*-S-*3-formamidosalicylate starter unit) and not the remaining genes (*antABCDE*) in the BGC (9). This also presumably indicates that starter unit biosynthesis is not rate limiting for antimycin production.

### Model for the regulation of antimycin biosynthesis

Our model for the regulation of antimycin biosynthesis is depicted in Fig. 6. Expression of the *ant* BGC is cross-activated by FscRI, a LuxR-family regulator, from the candicidin BGC, which activates expression of *antBA* and *antCDE* (10). This regulation in turn enables direct activation of the 3-FSA biosynthetic operons (*antGF* and *antHIJKLMNO*) by σ^AntA^. The expression of *antBA* and *antCDE* is down regulated following the onset of morphological differentiation, presumably because the ligand sensed by the FscRI PAS domain is no longer available (9–10). The cellular level of σ^AntA^ is antagonised by the ClpXP protease, for which it is a direct target, and is ultimately responsible for clearing residual σ^AntA^ when FscRI is inactivated following the onset of morphological differentiation (10). While ClpXP proteolytic control of transcription factor activity, and in particular that of ECF σ factor / anti- σ factors, has been shown previously (27, 28), (29–32); (33, 34); (35) it has thus far not been directly linked to the control of cluster-situated regulators of natural product biosynthesis. This finding provides a new lens through which to examine microbial signal transduction and the regulation of natural product biosynthesis in *Streptomyces* species. Understanding the diversity of regulatory strategies controlling the expression of these pathways is critical for the development of new tools for exploiting the ‘silent majority’ of biosynthetic pathways harbored by these organisms.

## Experimental procedures

### Growth media, strains, cosmids, plasmids, and other reagents

*Escherichia coli* strains were propagated on Lennox agar (LA) or broth (LB) (36, 37) and *Streptomyces albus* S4 strains were cultivated using LA, LB, and mannitol-soya flour (MS) agar or broth (36). Development of *clp* mutants was assessed on MS and ISP2 medium (36). Culture medium was supplemented with antibiotics as required at the following concentrations: apramycin, 50 μg ml^−1^; carbenicillin, 100 μg ml^−1^; chloramphenicol, 25 μg ml^−1^; hygromycin, 50 μg ml^−1^; kanamycin, 50 μg ml^−1^; nalidixic acid, 25 μg ml^−1^. *Streptomyces* strains were constructed by conjugal mating with *E. coli* ET12567 as previously described (36). Enzymes were purchased from New England Biolabs unless otherwise stated, and oligonucleotides were purchased from Integrated DNA Technologies, Inc. All of the strains, cosmids, and plasmids used in this study are described in Table S2, and all of the oligonucleotides used are provided in Table S3.

### Construction of plasmids

The insert for each plasmid generated in this study was prepared by PCR amplification with Q5 High-Fidelity DNA polymerase and oligonucleotides containing restriction sites. PCR-amplified inserts were restricted and cloned into the relevant plasmids cut with the same enzymes by standard molecular biology procedures. All clones were sequenced to verify the integrity of insert DNA. The restriction sites used for cloning are provided with the plasmid descriptions in Table S2.

### ChIP-sequencing and bioinformatics analyses

The *antA* coding sequence was amplified with RFS629 and RFS630, which contain KpnI and EcoRI restriction sites, respectively. The restricted PCR product was cloned into pSETNFLAG (10) digested with the same enzymes. The resulting plasmid was then restricted with NotI and EcoRI to release *ermE*\*p-3xFLAG-*antA*, which was subsequently cloned into pAU3-45 (38) digested with the same enzymes. pAU3-45-3xFLAG-*antA* was mobilised to an apramycin-marked Δ*antA* strain (9). Cultivation of the wild-type and Δ*antA*/pAUNFLAG-*antA* strains for ChIP-sequencing were performed exactly as described previously (10). The pure DNA resulting from immunoprecipitates from two biological replicates of wild-type and Δ*antA*/pAUNFLAG-*antA*, as well non-immunoprecipitated chromosomal DNA, were sequenced with the Illumina HiSeq3000 platform with 150-nucleotide paired-end reads by the University of Leeds Next Generation Sequencing Facility at the St. James Teaching Hospital NHS Trust. The resulting reads were analysed exactly as described previously (10). The graphic in Fig. 2 was generated using DeepTools computeMatrix and plotProfile functions (39).

### Construction of the *S. albus* S4 Δ*clpXclpP1clpP2* mutant strain

Deletion of *clpXclpP1clpP2* was carried out using RecET recombineering in *E. coli* as follows. The *clpXclpP1clpP2*-containing cosmid, cos117 was obtained by screening a previously constructed *S. albus* S4 Supercos1 cosmid library (8) by PCR using oligonucleotides PBB001 and PBB002. Cos117 was mutagenized as required using *E. coli* recombineering with strain GB05-red (40) and a deletion cassette. The deletion cassette was generated by PCR from paac-apr-oriT (41) and consisted of the apramycin resistance gene, *aac(3)IV* and a conjugal origin of transfer (*oriT*), which was flanked by ΦC31-*attL* and -*attR* sites for excision of the cassette. Oligonucleotides used to generate deletion cassettes included 39 nt of homology upstream or downstream of the target open reading frame(s) and are listed in Table S3. The resulting PCR product was digested with DpnI, gel purified and electroporated into arabinose-induced *E. coli* GB05-red harboring cos117. Transformants were screened for the presence of mutagenized cosmid by PCR using oligonucleotides listed in Table S3 and the integrity of the locus was verified by DNA sequencing. The mutagenized cosmid was electroporated into *E. coli* ET12567/pUZ8002 and mobilized to a strain of *S. albus* S4 harboring an entire antimycin BGC deletion (Δantall) (42) by conjugation as described (36). Transconjugants were screened for apramycin resistance and kanamycin sensitivity. The integrity of an apramycin-marked mutant was verified by PCR using the oligonucleotides listed in Table S3. The apramycin deletion cassette was subsequently excised from the chromosome by conjugal introduction of pUWLint31, which is a replicative plasmid with a temperature sensitive origin of replication that expresses the ΦC31 integrase required for removal of the cassette (41). Transconjugants were screened for loss of apramycin resistance and excision of the cassette was verified by polymorphic shift PCR and DNA sequencing of the product.

### Immunoblot analysis

Spores of the parental strain, *S. albus* Δantall and Δ*clpXclpP1clpP2* mutant harboring pPDA or pPDD were grown on MS agar (buffered with 50mM TES, pH 7.2) covered with cellophane discs. Protein extracts were prepared from mycelia collected at regular intervals during growth: 14h, 17h, 24h and 30h for Δantall and Δ*clpXclpP1clpP2* harboring 3xFLAG-AntA constructs; 17h, 20h, 23h and 30h for Δantall and Δ*clpXclpP1clpP2* harboring the 3xFLAG-FscRI construct. Protein extracts were generated as follows: 100 mg of cells were resuspended in 200 μl lysis buffer (50 mM sodium phosphate buffer, pH 7.0, 150 mM sodium chloride, 10 mg ml^−1^ lysozyme, cOmplete, Mini, EDTA-free protease inhibitors (Roche) and 100 mg of 0.1 mm glass beads (PowerLyzer®)) and lysed by vortexing for 30 min at 2000 pm, 37°C, with a subsequent incubation for another 30 min at 37°C. The obtained suspension was centrifuged for 20 min at 20,000 x *g* at 18°C. Thirty micrograms of the clarified protein extract were subjected to SDS-PAGE and then transferred to nitrocellulose membrane (pore size 0.2 μm) for Western blot analysis. The membrane was probed with mouse monoclonal ANTI-FLAG® M2-Peroxidase (HRP) antibody (Sigma), 1:10 000, and the signals were detected using Pierce™ 1-Step Ultra TMB Blotting Solution (Thermo Scientific).

### Protein purification and *in vitro* ClpXP proteolysis assays

The wild-type *antA* gene was PCR amplified and cloned into the AgeI and HindIII sites of the pET23b-SUMO vector, which harbors an N-terminal (His)_6_-SUMO tag (Wang *et al.* 2007). The plasmid for production of (His)_6_-SUMO-σ^AntA-DD^ was generated by site-directed mutagenesis (Agilent QuikChange) using primers listed in Table S3. (His)_6_-SUMO-σ^AntA^ and (His)_6_-SUMO-σ^AntA-DD^ were produced by *E. coli* Rosetta(DE3) (Novagen) grown in LB at 37 °C until OD_600_ 0.5, followed by induction with 0.4 mM IPTG and growth at 18 °C for 16 hours. Cells were resuspended in 50 mM sodium phosphate, pH 8, 1M NaCl, 20 mM imidazole, 10% glycerol, and 1 mM DTT and lysed by french press at 28 kpsi, followed by treatment with protease inhibitor cocktail set III, EDTA-free (Calbiochem) and benzonase (Millipore Sigma). (His)_6_-SUMO-σ^AntA^ and (His)_6_-SUMO-σ^AntA-DD^ proteins were purified by Ni-NTA affinity chromatography and Superdex-75 gel filtration and stored in 50 mM potassium phosphate, pH 6.8, 850 mM KCl, 10% glycerol, and 1 mM DTT. *E. coli* ClpX and ClpP proteins were purified as described previously (43, 44).

*In vitro* ClpXP proteolysis assays were performed at 30 °C by preincubating 0.3 μM ClpX_6_ and 0.8 μM ClpP_14_ with ATP regeneration system (4 mM ATP, 50 μg ml^−1^ creatine kinase, 5 mM creatine phosphate) in 25 mM HEPES-KOH, pH 7.5, 20 mM KCl, 5 mM MgCl2, 10% glycerol, 0.032% NP40, and 0.2 mM DTT and adding substrate to initiate the reactions. Samples of each reaction were taken at specific time points and stopped by addition of SDS-PAGE loading dye and boiling at 100 °C before loading on Tris-Glycine-SDS gels. Bands were visualized by staining with colloidal Coomassie G-250 and quantified by ImageQuant (GE Healthcare) after scanning by Typhoon FLA 9500 (GE Healthcare). The fraction (His)_6_-SUMO-σ^AntA^ remaining was calculated by dividing the (His)_6_-SUMO-σ^AntA^ density at a given time point by the density at time zero and normalized by ClpX density.

### Chemical analysis

*S. albus* S4 strains were cultivated atop a cellophane disc on MS agar at 30 °C for 7 days in triplicate. At the time of harvest, the cellophane disc containing mycelia was removed and the quantity of biomass was determined. Bacterial metabolites were extracted from both the mycelia and the ‘spent’ agar for 1 hr using 50 ml of ethyl acetate. Thirty millilitres of ethyl acetate were evaporated to dryness under reduced pressure and the resulting residue was resuspended in 100% methanol (300 μl). Immediately prior to LC-HRMS analysis, methanolic extracts were centrifuged at 16,000 × *g* in a microcentrifuge tube for 5 min to remove insoluble material. Only the supernatant (3 μl) was injected into a Bruker Maxis Impact TOF mass spectrometer equipped with a Dionext Ultimate 3000 HPLC as previously described (45). The peak area associated with the extracted ion chromatograms for antimycin A_1_, A_2_, A_3_ and A_4_ present in agar and mycelial extracts was determined and used to calculate the total antimycins produced for each replicate. These values were subsequently used to determine the arithmetic mean for total antimycin production for each strain. Statistical significance was assessed in MS Excel by a homoscedastic Student’s t-test with a two-tailed distribution.

## Data availability

The next-generation sequencing data obtained in this study are available under ArrayExpress accessions E-MTAB-7700 and E-MTAB-5122.

## Acknowledgements

We thank Matt Hutchings, Paul Hoskisson, Kenneth McDowall and Alex O’Neill for helpful discussion concerning this manuscript.

## Author contributions

BB, SK and AF performed experiments, interpreted data and wrote sections of the manuscript; TAB wrote sections of the manuscript; RFS designed the study, performed experiments and wrote the manuscript.

