## Supplementary information for "Regulation of antimycin biosynthesis is controlled by the ClpXP protease"




**Running title:** Regulation of antimycin biosynthesis is controlled by the ClpXP

**Table of contents:**

**Fig. S1.** MUSCLE alignment of 71 $\sigma^{AntA}$ orthologues
**Fig. S2.** Gel filtration of purified $(His)_6$-SUMO-$\sigma^{AntA}$ proteins
**Fig. S3.** Sporulation of $\Delta$*clpXclpP1clpP2* and parental strains
**Fig. S4.** Uncropped western blots
**Table S1.** Relative abundance of antimycins produced by $\Delta$*clpXclpP1clpXP2* and the parental strain
**Table S2.** Bacterial strains, cosmids and plasmids used in this study
**Table S3.** Oligonucleotides used in this study
**References**

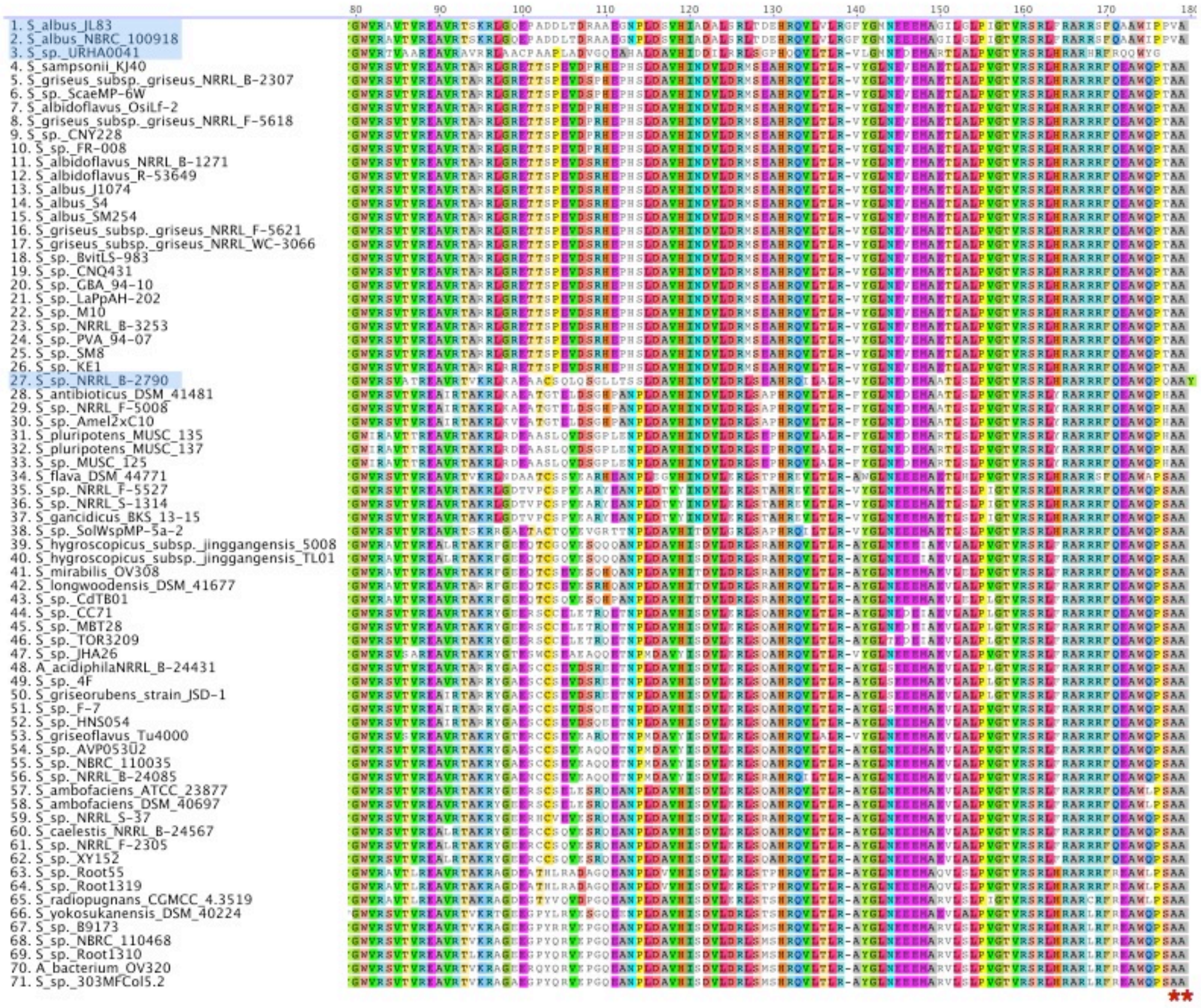


**Fig. S1. MUSCLE alignment of 71 σ$^{AntA}$ orthologues.** Only alignment of amino acid residues 80-180 are shown. Red asterisks indicate the C-terminal Ala-Ala motif conserved in 67 of 71 orthologues. Blue shading indicates of taxa names indicate those that do not possess the motif. The orthologue from *Streptomyces* sp. NRRL B-2790 terminates in Ala-Ala-Tyr; the inclusion of a terminal Tyr residue is likely an artefact caused by a poor genome sequence quality (>4000 contigs). The remaining three orthologues lacking the di-alanine motif are *Streptoacidiphilus albus* strains (terminating in Val-Ala) and *Streptomyces* sp. URHA-0041 (Tyr-Gly). Interestingly, these three taxa are progenitors within the proposed evolutionary trajectory of the *ant* BGC (1) and the lack of the motif may mark a point where Clp-protease degradation of σ$^{AntA}$ had yet to evolve.

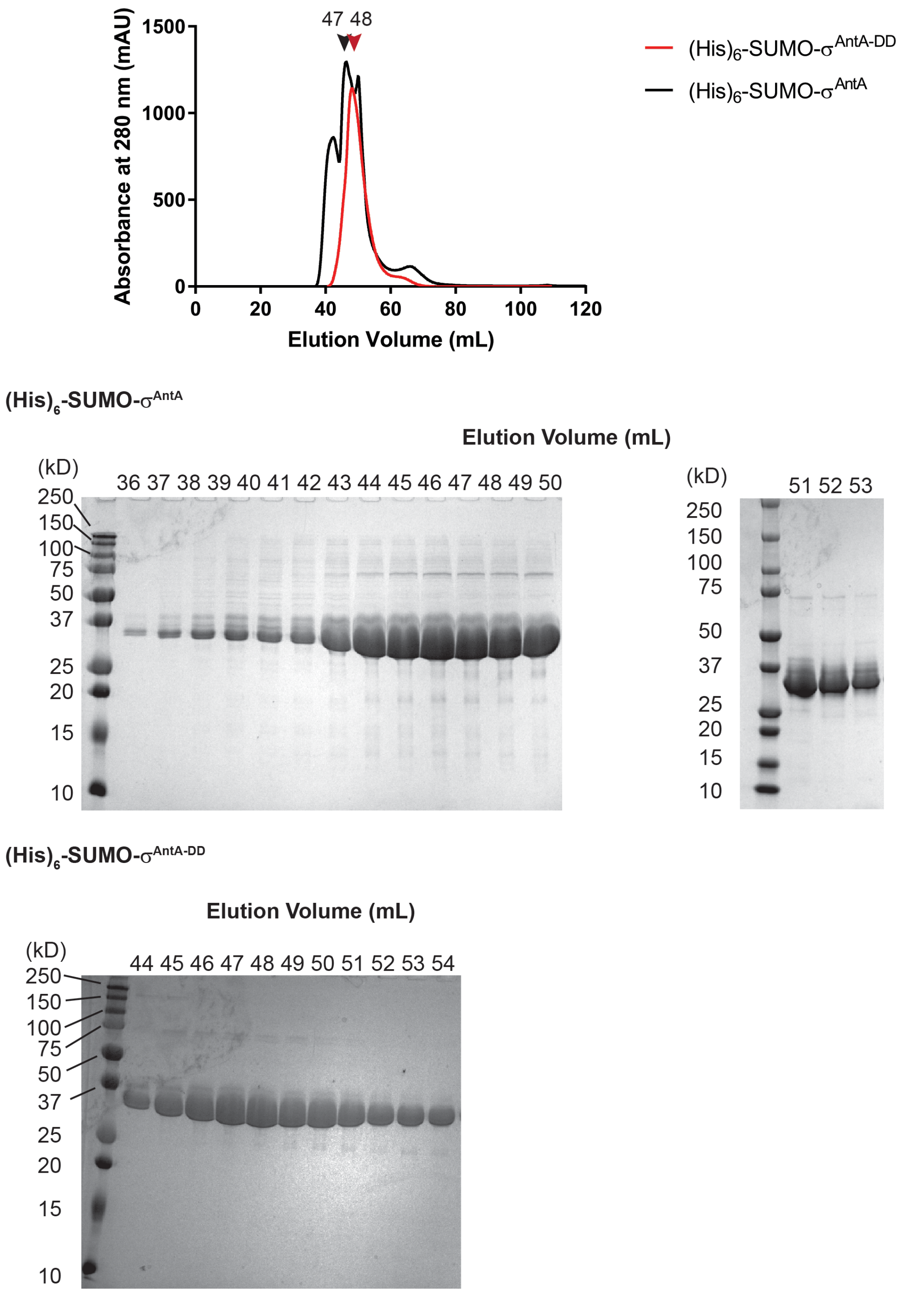


**Fig. S2. Gel filtration of purified (His)$_6$-SUMO-$\sigma^{AntA}$ proteins.** Gel filtration chromatograms showing the elution profiles of purified (His)$_6$-SUMO-$\sigma^{AntA}$ and (His)$_6$-SUMO-$\sigma^{AntA\text{-}DD}$ proteins (upper panel) and associated SDS-PAGE analyses (lower panels). The chromatogram of (His)$_6$-SUMO-$\sigma^{AntA}$ differs slightly from (His)$_6$-SUMO-$\sigma^{AntA\text{-}DD}$ because gel filtration of (His)$_6$-SUMO-$\sigma^{AntA}$ protein was performed in 50 mM HEPES-KOH pH 7.5, 150 mM KCl, 10% glycerol, and 1 mM DTT and later dialyzed into the storage buffer, whereas gel filtration of (His)$_6$-SUMO-$\sigma^{AntA\text{-}DD}$ was performed in the storage buffer.

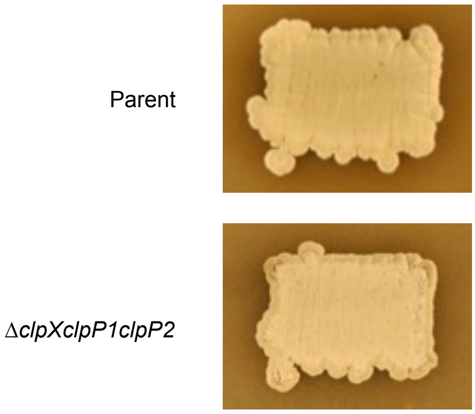


**Fig. S3. Sporulation of *S. albus* S4 *clp* mutants.** Photographs were taken after 6 days of growth on the indicated medium. The Δ*clpXclpP1*Δ*clpP2* strain undergoes a normal developmental cycle, however sporulation is less robust on MS agar media compared to the parental strain.

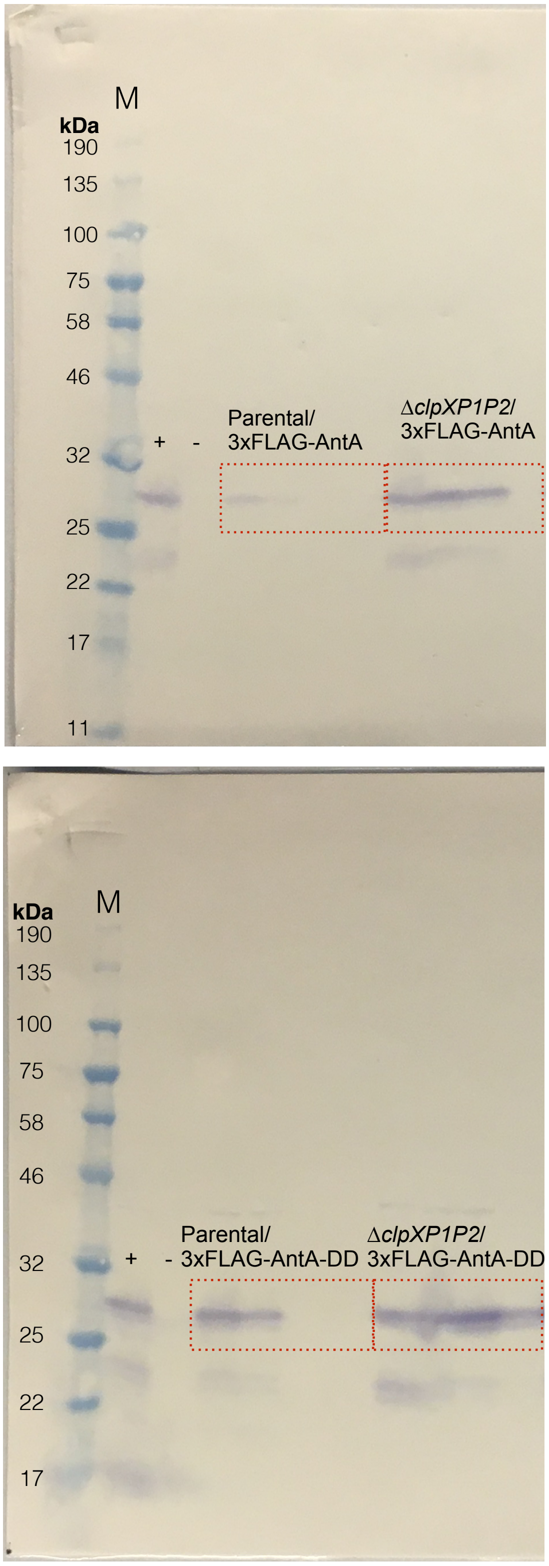


**Fig. S4. Uncropped western blots.** Shown are the original western blots depicted in Fig. 4. Red boxes indicate cropped sections for the strains indicated. The Mw of the protein marker (M) is shown. Plus and minus symbols on each blot indicate positive (Δ*clpXclpP1clpP2/*pPDD) and negative controls (parental strain Δantall), respectively.

**Table S1. LCMS quantification of antimycin production**

| | Total antimycin peak area (arbitrary units) | | | | |
|---|---|---|---|---|---|
| ***S. albus* S4 strain** | **Replicate 1** | **Replicate 2** | **Replicate 3** | **Average** | **SD*** |
| Parental (Δantall) *attB* ΦC31 cos213 | 17.24 | 15.3 | 17.24 | 16.59 | 1.12 |
| ΔantallΔ*clpXclpP1clpP2 attB* ΦC31 cos213 | 18.71 | 13.11 | 14.88 | 15.57 | 2.86 |

* SD = Standard Deviation

**Table S2. Bacterial strains, cosmids and plasmids used in this study**

| Name | Description | Reference or source |
|---|---|---|
| ***Streptomyces* strains** | | |
| *S. albus* S4 | Wild type *Streptomyces albus* S4 strain | (2) |
| *S. ambofaciens* ATCC 23877 | *Streptomyces ambofaciens* wild-type strain | ATCC |
| Δ*antA* | *S. albus* S4 harbouring an *antA* deletion; $Apr^R$ | (3) |
| Δ*antA*/pAU3-45-3xFLAG-*antA* | Δ*antA* strain harbouring pAU3-45-3xFLAG-*antA* at the ΦC31 *attB* site; $Apr^R$, $Tsp^R$ | This study |
| Δ*antall* | *S. albus* S4 harbouring an unmarked deletion removing the entire antimycin BGC | (4) |
| Δ*clpXclpP1clpP2* | Δ*antall* strain with an unmarked mutation deletion removing the *clpX, clpP1, clpP2* operon | This study |
| ***Escherichia coli*** | | |
| XL10-Gold | General cloning host | Stratagene |
| ET12567 | Non-methylating host for transfer of DNA into *Streptomyces* spp. (*dam, dcm, hsdM*); $Cam^R$ | (5) |
| GB05-red | Host for RecET recombination | (6) |
| Rosetta(DE3) | Host for heterologous protein production | Novagene |
| **Cosmids** | | |
| cos117 | Supercos1 derivative containing the *clpP1P2* and *clpX* genes; $Carb^R$, $Kan^R$ | This study |
| c117Δ*clpXP*::*aac(3)IV* | The c117-derivative with the *clpP1P2* and *clpX* replaced by the disruption cassette from paac+oriT; $Carb^R$, $Kan^R$, $Apr^R$ | This study |
| **Plasmids** | | |
| patt-saac-oriT | PCR template for *aac3(IV)* oriT cassette used in REDIRECT PCR targeting system; $Apr^R$, $Amp^R$ | (7) |
| pAU3-45 | pSET152 derivative, integrates into *φ*C31 attachment site; $Apr^R$ $Tsp^R$ | (8) |
| pAU3-45-3xFLAG-*antA* | pAU3-45 derivative containing the *ermE**p-3xFLAG-*antA* cloned into the NotI and EcoRI sites; $Apr^R$, $Tsp^R$ | This study |
| pET23b-His-SUMO | pET23b with a SUMO tag cloned into the NheI-AgeI sites; $Carb^R$ | (9) |
| pET23b-His-SUMO-*antA* | pET23b-His-SUMO derivative with the wild-type *antA* gene from *Streptomyces ambofaciens* ATCC 23877 cloned into the AgeI-HindIII sites; $Carb^R$ | This study |
| pET23b-His-SUMO-*antA-DD* | pET23b-His-SUMO-*antA* derivative harbouring with point mutations changing the AntA C-terminal AlaAla to AspAsp; $Carb^R$ | This study |
| pPDA | pSETNFLAG derivative harboring *antA* cloned into KpnI-EcoRI sites; $Apr^R$ | This study |
| pPDD | pSETNFLAG derivative harbouring *antA* encoding A172D and A173D mutations cloned into KpnI-EcoRI sites; $Apr^R$ | This study |
| pSET152 | *E. coli – Streptomyces* integrative shuttle vector, integrates into the ΦC31 attachment site; $Apr^R$ | (10) |
| pSETNFLAG | pSET152 derivative with an *ermE**p cloned into the EcoRV-EcoRI sites; and an N-terminal 3xFLAG tag and multi-cloning site cloned into the NdeI-KpnI sites; $Apr^R$ | (11) |
| pUWLint31 | *Streptomyces* vector for the expression of the ΦC31 integrase; pSG5 temperature sensitive *ori*; $Tsp^R$ | (7) |
| pUZ8002 | Encodes conjugation machinery for mobilization of plasmids from *E. coli* to *Streptomyces*; $Kan^R$ | (5) |

Cam – chloramphenicol; Carb – carbenicillin; Kan – kanamycin; Apr – apramycin; Spr – spectinomycin; Hyg – hygromycin, Tsp - thiostrepton

**Table S3. Oligonucleotides used in this study**

| Name | Sequence (5’-3’)* | Description |
|---|---|---|
| PBB001 | gtggcacgcatcggtgacggc | PCR: to identify the *clpX*-containing cosmid |
| PBB002 | tcaggccgacttctgctcgcc | |
| PBB003 | gtgacgaatctgatgccctcc | PCR: to confirm presence of the *clpP1P2* operon on the cosmid |
| PBB004 | tcagactccggcgttgagctt | |
| PBB034 | agacggcccggcgccgtcgtaagacgagcaggtggatacttccggggatccgtcgaccc | PCR: *clpXP*/*aac3(IV)+oriT* recombineering cassette |
| PBB035 | ggtggggcccttccgcgtgcgtctgccgggtgccggccctgtaggctggagctgcttcg | |
| PBB015 | ccaaacggcgggcgcggaccc | PCR: to confirm the *clpXP* deletion |
| PBB018 | gacggaagggccccaccgcgc | |
| RFS629 | tatata*ggtacc*aacaccgcgcacgaactgccc | PCR: to amplify $\sigma^{AntA}$. Contains a KpnI site |
| RFS630 | tatata*gaattc*tcaggcggcggtgggctgcc | PCR: to amplify $\sigma^{AntA}$. Contains an EcoRI site |
| RFS663 | tatata*gaattc*tcagtcgtcggtgggctgc | PCR: to amplify $\sigma^{AntA}$, encodes A172D and A173D mutations. Contains an EcoRI site |
| SK221 | cgc*aagctt*tcacgccgcc | PCR: to amplify $\sigma^{AntA}$. Contains a HindIII site |
| SK222 | tat*accggt*ggttccaccgtcagcgaactcc | PCR: to amplify $\sigma^{AntA}$. Contains an AgeI site |
| SK232 | ggcatggctgccctcggatgattga*aagcttgcggccgc* | PCR: to mutagenize $\sigma^{AntA}$. Contains HindIII and NotI sites |
| SK233 | *gcggccgcaagctt*tcaatcatccgagggcagccatgcc | PCR: to mutagenize $\sigma^{AntA}$. Contains HindIII and NotI sites |

*Restriction sites are indicated by italics and non-homologous sequences are underlined
